# Helicobacter pylori cancer associated CagA protein drives intestinal metaplastic transition in human gastric organoids

**DOI:** 10.1101/2023.11.22.567682

**Authors:** Mar Reines, Meike Soerensen, Hilmar Berger, Mihir Patel, Philipp Schlärmann, Thomas F. Meyer

**Affiliations:** Max Planck Institute for Infection Biology, Department of Molecular Biology, Berlin, Germany; Institute of Clinical Molecular Biology, Christian Albrecht’s University of Kiel, Germany

**Keywords:** Organoids, bacterial infection, type 4 secretion, STAT3 activation, CDX2, epithelial trans-differentiation

## Abstract

Gastric intestinal metaplasia (GIM) constitutes a pre-neoplastic stage in the development of stomach cancer. While strong evidence points to a role of infection with CagA positive *Helicobacter pylori* in the development of GIM, currently available experimental models have not provided mechanistic clues on this association. Here, we ectopically expressed the *H. pylori* CagA protein in human gastric organoids derived from normal, primary epithelial cells. Native CagA protein was produced and rapidly processed to yield a tyrosine-phosphorylated C-terminal fragment of ∼35 kDa. It led to an activation of the STAT3 pathway and aberrant elevation of CDX2 expression, a marker of intestinal type of cells, as well as other intestinal markers. Thus, CagA drives re-programming of gastric cells towards an intestinal-like phenotype, towards GIM. In summary, we describe a cooperative mechanism of CagA-induced STAT3 signaling and intestinal-like trans-differentiation, promoting a pre-neoplastic state. Our model provides mechanistic evidence for a direct role of CagA in driving premalignancy in gastric pathogenesis.

## Introduction

Gastric cancer is the fourth leading cause of cancer-related mortality in the world ^1,2^. Typically, it progresses through a defined sequence of pathological stages, starting with chronic active gastritis, followed by atrophic gastritis, gastric intestinal metaplasia (GIM), and finally adenocarcinoma ^3,4^. The main risk factor of this cascade of pathologic events and gastric cancer development is infection with the Gram-negative bacterium *Helicobacter pylori* ^5,6^.

*H. pylori* colonizes the stomach of ∼50% of the world’s population ^7^. Although its prevalence has decreased in Western countries, it is still responsible for more than half a million of gastric cancer deaths world-wide, annually. The pathogenesis and severity of *H. pylori* infections are strongly influenced by whether the infecting strain expresses the bacterial CagA protein. It is encoded by the cag pathogenicity island cagPAI and translocated into gastric epithelial cells via the type IV secretion system (T4SS) ^8^ along with a release of ADP-heptose, driving inflammation ^9,10^. Upon translocation, CagA undergoes rapid tyrosine phosphorylation by members of the SRC family and c-Abl kinases ^11–14^. Phosphorylated CagA has been shown to deregulate several host signalling pathways, thus disturbing gastric epithelial homeostasis^15,16^.

Several animal models have shown a link between CagA and cancerogenesis. Mongolian gerbils infected with a CagA-positive *H. pylori* strain develop gastric carcinoma ^17^. Moreover, in transgenic mice, ectopic CagA expression leads to a spontaneous development of gastric cancer ^18^ and in a zebrafish model, CagA overexpression enhances intestinal carcinogenesis ^19^. Additionally, recent studies have also used human models to investigate the mechanisms by which CagA exerts oncogenic effects. In human gastric organoids, the interaction between CagA and the apoptosis-stimulating protein of p53 2 (ASPP2) promotes the survival of CagA-positive *H. pylori* in the organoid lumen due loss of cell polarity ^20^. Moreover, CagA-positive *H. pylori* infection inhibits p53 stabilization in the nucleus, promoting genetic instability and fast progression towards carcinogenesis ^21^.

Interestingly, CagA also appears to be associated with the pre-cancerous stage of GIM ^4^. GIM is histopathologically characterized by the replacement of gastric cells with intestinal-like cells, including enterocytes and goblet cells ^22^. Apart from a variety of genetic and epigenetic alterations ^23^, GIM is characterised by the abnormal expression of the transcription factor CDX2, which is normally expressed exclusively in intestinal epithelial cells ^24–30^. CDX2 is essential for intestinal differentiation by maintaining the expression of intestinal genes, such as the goblet cell marker mucin MUC2 and TFF3 ^31,32^. Several studies have linked infections with CagA-positive *H. pylori* strains to increased CDX2 expression in gastritis and GIM biopsies ^3,22,33,34^, however, a direct role of CagA in that process has not been revealed. Furthermore, GIM is thought to emerge in the context of elevated inflammation and regeneration of the stomach mucosa^35^, with the STAT3 pathway being one of the most activated oncogenic pathways in the pre-cancerous development ^36–38^.

In the present study, to assess the role of CagA in cellular transformation, we used normal human gastric organoids to ectopically express CagA. In our system, CagA was functionally processed to yield a 35 kDa C-terminal tyrosine-phosphorylated protein. It led to constitutive activation of the STAT3 pathway and upregulation of CDX2, as well as downstream intestinal markers, thus driving the re-programming of gastric cells towards intestinal-like cells. These observations highlight the direct link between CagA-induced trans-differentiation and activation of the pro-tumorigenic STAT3 pathway, revealing insights into the molecular signaling induced by *H. pylori* to drive cellular transformation.

## Results

### In human gastric organoids, ectopically expressed CagA is processed into a 35 kDa C-terminal fragment subjected to tyrosine phosphorylation

In order to analyze the causative role of CagA in gastric pathology, we chose to express this effector protein in organoids derived from healthy human gastric epithelial cells^39–41^. In contrast to transformed human cell lines, the cellular physiology and signaling pathways in organoids remain unaltered, making this model more appropriate for studying pathology and early events of transformation. Cultured gastric organoids were transduced with lentivirus bearing *gfp* and *H. pylori cagA* under the control of an EF promoter (Fig. 1A). As a control, gastric organoids were transduced with lentivirus containing only *gfp* under a CMV promoter (Fig. S1A). Ten days post-transduction, stably transduced cells were selected by FACS sorting of single GFP^+^ cells derived from the transgenic organoid cultures. Transgenic CagA-positive stem cells grown in expansion medium were able to re-generate organoids within 14 days after FACS. Furthermore, organoids kept expanding as a homogenously CagA-positive culture (Fig. 1B and Fig. S1A). No major structural changes were observed.

**Fig. 1.**
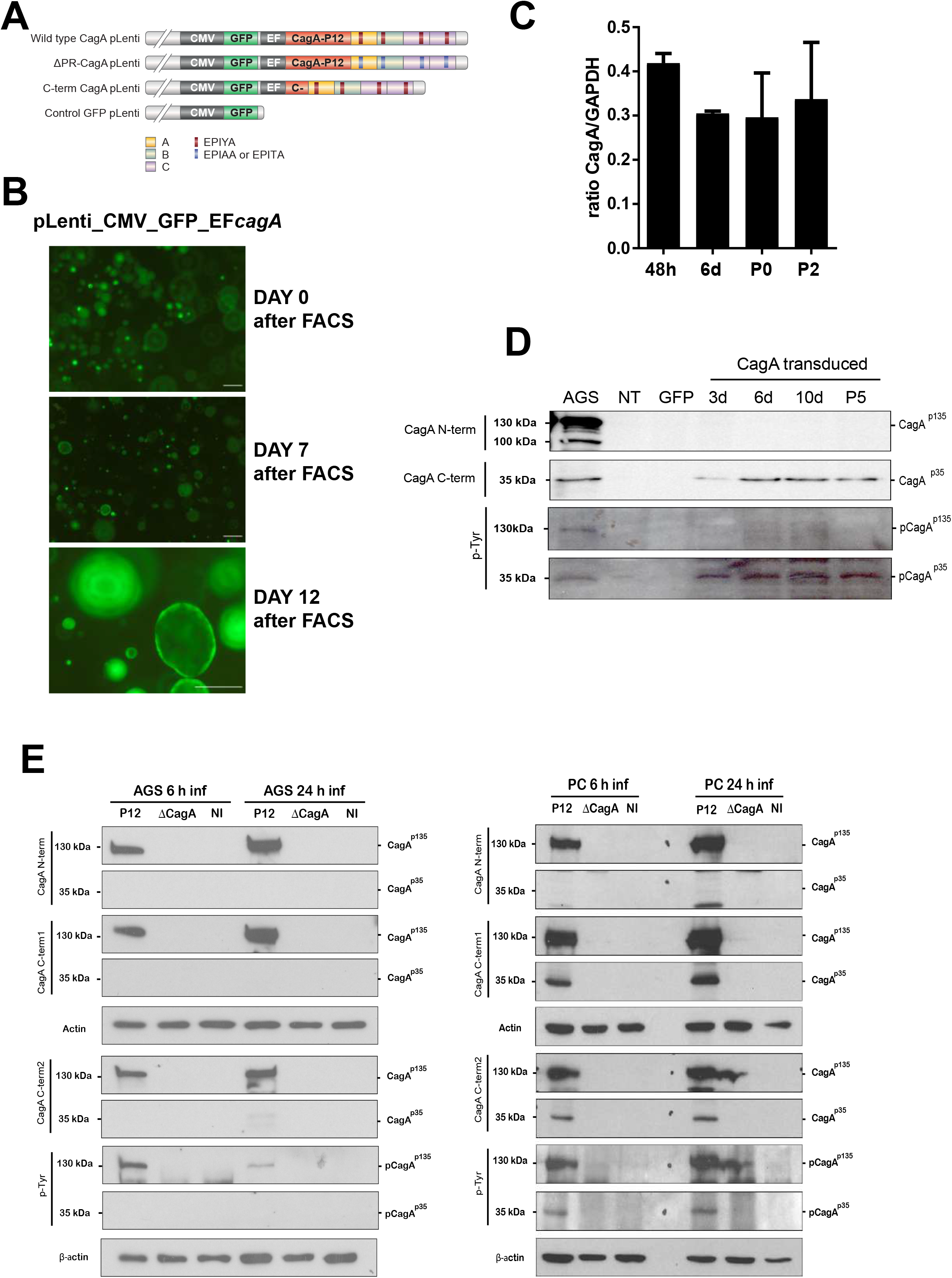
*H. pylori* CagA is processed and phosphorylated in human gastric organoids. Gastric organoids were transduced with lentivirus MOI 0.5, positively selected with FACS, and expanded as homogenous cultures. (A) Lentivirus constructs expressing CagA and CagA mutants under the control of the EF1alpha promoter. The *H. pylori* CagAs used in this study: The western-type CagA characterized by the presence of the EPIYA-C segment duplicated in tandem (Wild type CagA); a phospho-resistant CagA with 4 mutated tyrosine phosphorylation sites, where tyrosine is exchanged to alanine (EPIYA to EPIAA) (ΔPR-CagA); the C-terminal part of CagA (67 KDa) (C-term). As a control of efficient transduction a construct only expressing GFP under CMV promoter was used (Control GFP) (B) Successful lentiviral transduction in human gastric organoids was monitored by GFP expression under the control of the CMV promoter. Representative micrographs of human transgenic gastric organoids expressing GFP and CagA, FACS sorted as single cells (Day 0) and cultured (Day 5 and Day 12). Scale bars: 500 µm. n=7. (C) Time-course by RT-qPCR of CagA expression in human transgenic gastric organoids at 48 hours,6 days post lentiviral transduction, and in passage 0 (P0) and passage 2 (P2) after FACS. Values of mRNA levels were normalized against GAPDH. Data represent mean ± SD of more than five independent biological experiments. (D) Western blot analysis of CagA expression and phosphorylation in human transgenic gastric organoids using antibodies against the N-terminal part (b-300, N-term) and the C-terminal part (bk-20, C-term) of CagA. Phosphorylated CagA was detected using the pTyr99 antibody. Indicated in the blot: full-length CagA (CagA^p135^), phosphorylated full-length CagA (pCagA^p135^), C-term cagA (CagA^p35^), phosphorylated C-term CagA (pCagA^p35^). The results are representative of three independent experiments. AGS cells transduced with wild type *cagA-* lentivirus and human gastri organoids transduced with Control GFP were used as a control. (E) Western blot analysis of CagA expression and phosphorylation in AGS and 2D human gastric organoids infected with *H. pylori* P12 MOI 100 for 6 h and 24 h. Different anti-CagA antibodies were used to specifically detect the whole protein (CagA^p135^), the N-terminal part (CagA^p100^), and the C-terminal part (CagA^p35^). Phosphorylated CagA was detected using the pTyr99 antibody (CagA^p135^, CagA^p35^). The results are representative of three independent experiments.

To evaluate the CagA expression levels in the transgenic organoids, we first performed a time-course RT-qPCR. The expression of CagA mRNA in primary cells was stable over several passages (Fig. 1C). We next examined the transgenic organoids for CagA protein expression but could not detect the 130-135 kDa protein ^8,42^ in contrast to the positive control of CagA-transduced AGS cells (CagA^p135^) (Fig. 1D). Previous studies in our group ^43–46^ have shown that in infected phagocytic-like cell lines, translocated CagA can be processed into three different protein products: a full-protein of 135 kDa, an N-terminal part of 100 kDa and a C-terminal part of 35 kDa. These CagA products have also been detected in infected B cell lines ^47^. Thus, we sought to assess if this was also the case for CagA expressing gastric organoids. We employed the bk-20 antibody, which recognizes the C-terminal part of CagA, in order to detect secondary products of CagA breakage. Indeed, we detected a truncated CagA protein of ∼35 kDa in the transgenic organoids as well as in the transduced AGS cells (CagA^p35^, Fig. 1D). Our group has characterized this product as the C-terminal part of CagA, which contains the EPIYA motifs phosphorylated by Src family kinases (SFKs) and c-Abl kinase ^12,13^. Thus, we employed an anti-pTyr99 antibody to determine the phosphorylation status of the protein. CagA was confirmed as a tyrosine-phosphorylated 35 KDa product (pCagA^p35^, Fig. 1D). Finally, we transduced human gastric organoids with lentivirus containing HA-tagged C-terminal CagA. Using an anti-HA antibody, we verified the existence of a 35 KDa protein that corresponded to that detected by the anti-C-terminal CagA antibody (data not shown).

Upon translocation, CagA size modification and phosphorylation patterns differ between different cell types. Thus, we infected a 2-D organoid-derived monolayer with *H. pylori* P12, as this model provides ideal conditions to study infection mechanisms *in vitro* ^39^. In contrast to AGS cells, in these human primary gastric cells CagA was rapidly processed into a 100 kDa N-terminal (CagA^p100^) and tyrosine-phosphorylated 35 kDa C-terminal (CagA^p35^), in addition to the full-length CagA (CagA^p135^) (Fig. 1E). Interestingly, in human gastric primary cells, the fragmentation pattern of both *H. pylori* P12-translocated CagA and ectopically expressed CagA was similar. This phenomenon indicates that CagA undergoes cleavage processing into the functional protein in human gastric organoid cells. However, in organoids ectopically expressing CagA, the full CagA protein and the N-terminal part were not detected, neither by using the anti-N-terminal (b-300) antibody nor by using the anti-C-terminal (bk-20) antibody. The protease activity involved has yet to be identified.

### Gene expression profile in CagA-positive organoids

To assess the global cellular responses to ectopic CagA expression in human gastric organoids, we performed microarray analysis of FACS-sorted homogenous CagA-positive vs. control gastric organoids at passage 1 and passage 7 (Fig. S1B). Interestingly, stomach and intestinal-specifc markers were differentially regulated. The expression of intestinal differentiation markers (e.g., CDX2, CDX1, TFF3, and TFF1) was upregulated in CagA-positive gastric organoids compared to controls. Conversely, gastric differentiation markers (e.g., SOX2, and TFF2) were downregulated, indicating that CagA expression in gastric organoids modified cell identity by driving trans-differentiation towards intestinal-like organoids (Fig. 2A). Moreover, gene set enrichment analysis (GSEA) revealed a significant enrichment of the GIM signature ^48^ in CagA-positive gastric organoids compared to non-transduced controls (Fig. 2B). Furthermore, tissue-specific gene sets of the intestine showed significant positive enrichment in CagA-positive gastric organoids compared to controls (Fig. 2C and Fig. S2A). Altogether, these data suggest a role for CagA in promoting a GIM phenotype in gastric organoids *in vitro*.

**Fig. 2.**
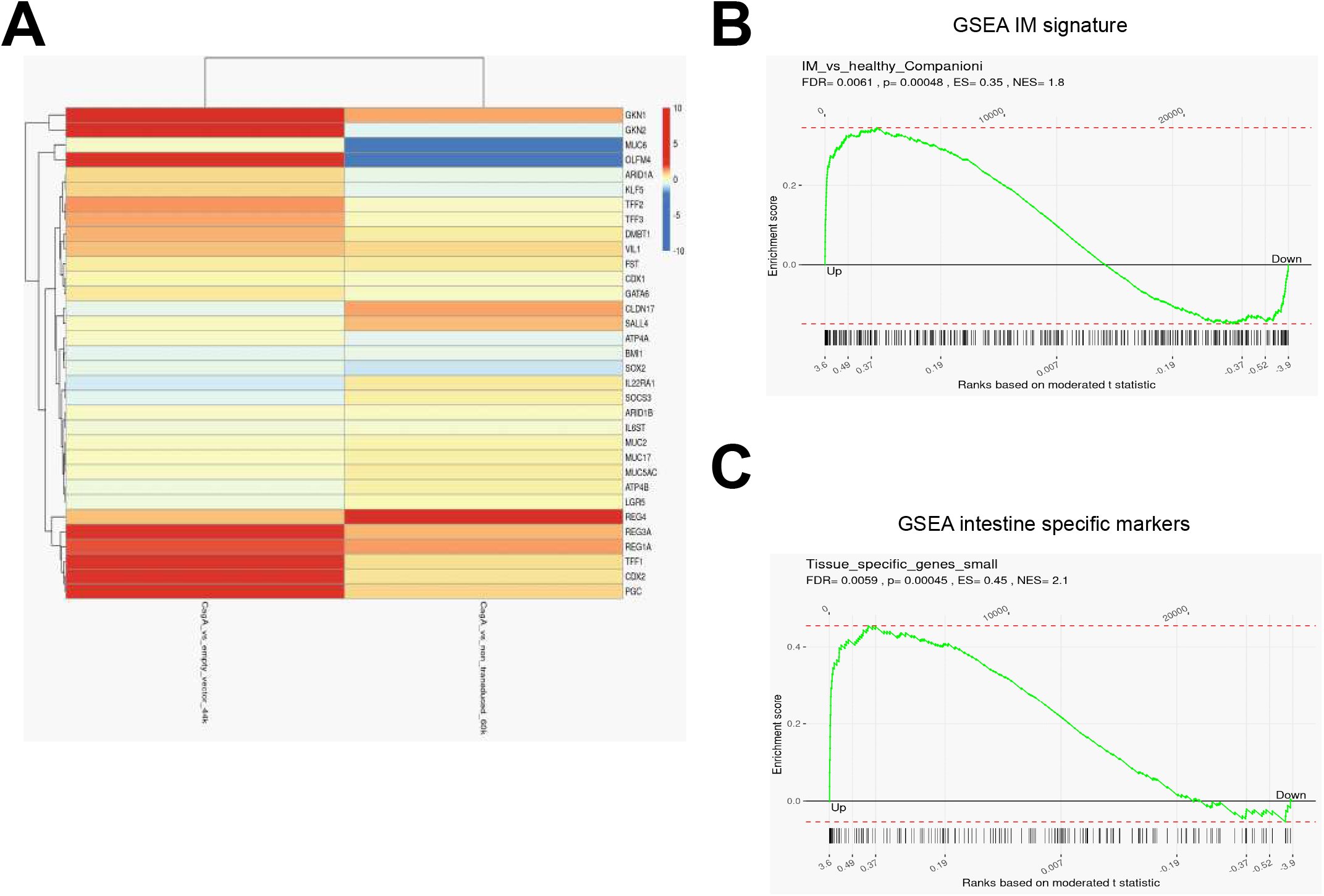
Microarray analysis of CagA-regulated genes. Healthy organoids were transduced with CMV*gfp*_EF*cagA* or an empty vector as a control, FACS sorted for GFP-positive cells and the homogenous population was expanded. We performed microarray analysis comparing CagA-positive organoids with control organoids in passage 0 (P0) and passage 7 (P7). (A) Heat map indicating the expression changes of selected gastric-specific and intestine-specific genes as determined by microarray analysis highlighted in the global overview. Data are mean-centered and averaged expression values for each replicate (n = 2). (B) Gene set enrichment analysis of the microarray results based on intestinal metaplasia marker genes ^48^. Genes are ordered according to their differential expression in CagA-positive organoids vs. control gastric organoids (left to right). Intestinal metaplasia marker genes are marked by black bars below the plot. The plotline indicates the running enrichment score. (C) Gene set enrichment analysis of the microarray results based on small intestine-specific markers (from Protein Atlas). Genes are ordered according to their differential expression in CagA-positive organoids vs. control gastric organoids (left to right). Tissue-specific small intestine genes are marked by black bars below the plot. The plotline indicates the running enrichment score.

### CDX2 drives intestinal-like differentiation in gastric organoids

To validate the microarray data, we used RT-qPCR to confirm the expression of CDX2, a master transcription factor of intestinal differentiation markers. Transgenic organoids were harvested at 48 h, 6 days, and 8 days post-lentivirus transduction, and RNA was isolated. RNA from organoids transduced with an empty vector served as control. CDX2 mRNA was strongly upregulated in CagA-positive organoids compared to controls, even 6 and 8 days after CagA transduction (Fig. 3A).

**Fig. 3.**
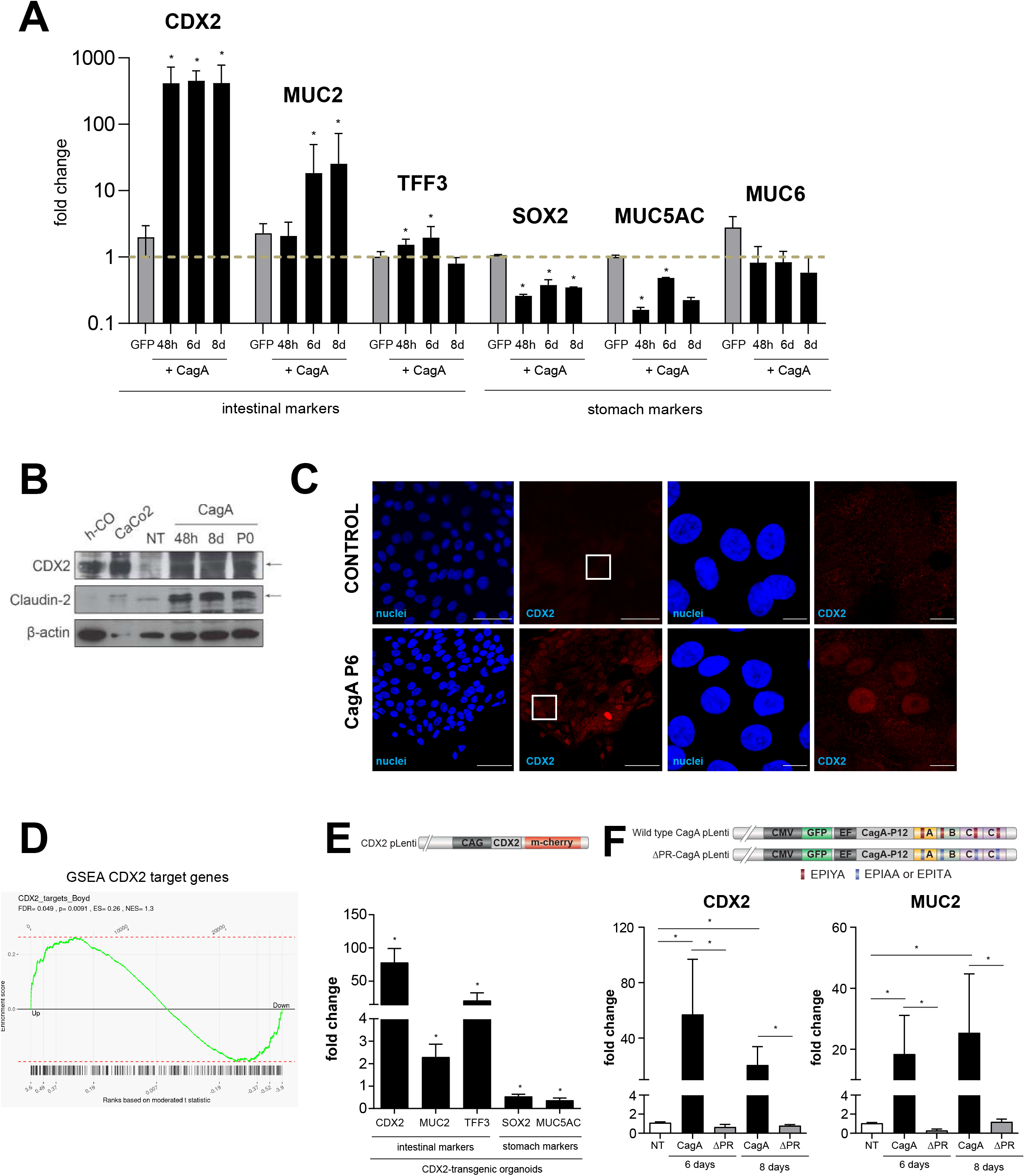
CagA expression in gastric organoids induces intestinal differentiation. (A) Gene expression analysis by RT-qPCR of the intestine-specific markers CDX2 and MUC2, and the stomach-specific markers SOX2, MUC5AC and MUC6 in CagA-positive organoids (CagA) and Control GFP organoids (GFP) at 48 h, 6 and 8 days post-transduction. Values of mRNA levels were normalized against GAPDH expression and further normalized against non-transduced (NT) cells (represented by the grid line at 1). Data represent mean ± SD of at least three independent biological experiments. Data is expressed as log10 (number format antilog). (B) Western blotting analysis of CDX2 and the CDX2 target Claudin 2 expression in CagA-positive organoids 48 h and 8 days post-transduction, and at P0 after FACS. Human primary colon epithelial cells (h-CO) and colon cancer cell line (CaCo2) were used as control. β-actin was used as a loading control. (C) Analysis of CDX2 translocation to the nucleus (red) by confocal IF, maximum projection. Cells were transduced with *cagA*-lentivirus, FACS sorted, and passaged until P6. Scale bar=10 µm (right panel: inset with detailed nucleus, scale bar= 500 µm) (D) Gene set enrichment analysis of the microarray results based on the CDX2 transcription factor target genes ^58^. Genes are ordered according to their differential expression in CagA-positive organoids vs. control gastric organoids (left to right). CDX2 transcription factor target genes are marked by black bars below the plot. The plotline indicates the running enrichment score. (E) Gene expression analysis by RT-qPCR of intestinal differentiation markers (CDX2, TFF3 and MUC2) and gastric differentiation markers (SOCX2 and MUC5AC) in gastric organoids overexpressing CDX2. Values of mRNA levels were normalized against GAPDH expression and further normalized against non-transduced (NT) cells. Data represent mean ± SD of three independent biological experiments. (F) Gene expression analysis of CDX2 and MUC2 by RT-qPCR in CagA-positive organoids (CagA) and phospho-resistant mutant CagA-positive (ΔPR-CagA) 6 days and 8 days post-transduction. Values of mRNA levels were normalized against GAPDH expression. Data represent mean ± SD of three independent biological experiments.

Next, we analysed expression levels of the transcription factor SOX2 in the CagA-positive organoids, which plays an important role in gastric differentiation during development ^49,50^. SOX2 expression is progressively lost in GIM and gastric adenocarcinoma ^51–54^. Furthermore, inverse expression levels of SOX2 and CDX2 are observed in the stomach mucosa during gastric carcinogenesis ^55^. Indeed, SOX2 mRNA levels were down-regulated in CagA-positive organoids (Fig. 3A).

We then looked at different mucins, as GIM of the gastric mucosa is linked to changes in the mucin profile. Mucin 2, a specific marker for intestinal differentiation ^26^, was upregulated in CagA-positive organoids compared to controls (Fig. 3A). Muc5AC, key mucin of the pit mucous cells, was down-regulated. However, Muc6, key mucin of normal gland mucous cells of the stomach, was not regulated in CagA-positive organoids (Fig. 3A).

Owing to the important role of CDX2 in directing and maintaining an intestinal phenotype, we analyzed CDX2 protein expression in CagA-positive organoids in detail using Western immunoblotting. We detected increased CDX2 protein at 48 h, 8 days, and at passage 0 (i.e., after FACS of Cag-A-GFP^+^ single cells, Fig. S1B) (Fig. 3B). Claudin-2, an integral transmembrane protein of the tight junctions, is a target gene controlled by CDX2 ^34,56,57^. At 48 h, 8 d, and passage 0, Claudin-2 was upregulated in CagA-positive organoids together with the upregulation of CDX2 (Fig. 3B). CDX2 acts as a transcription factor upon translocation to the nucleus. Immunofluorescence analysis of CagA-positive organoids indicated nuclear localization of CDX2 even after 6 passages (Fig. 3C).

GSEA of the transcriptome data of the CagA-positive organoids revealed a significant enrichment of CDX2 target genes ^58,59^ (Fig. 3D). Thus, we wondered if CDX2 could be sufficient to transform normal human gastric organoids towards cells with an intestinal-like identity. We transduced dissociated gastric organoids with a lentivirus expressing CDX2 controlled by the CAG promoter, coupled to an mCherry reporter gene (Fig. S3A). At 8 days post-transduction, CDX2-positive organoids overexpressed CDX2 and the intestinal markers MUC2 and TFF3 compared to control organoids (Fig. 3E). Moreover, CDX2-positive organoids downregulated the gastric markers SOX2 and MUC5AC (Fig. 3E). Expression of intestinal markers and downregulation of gastric markers indicates a degree of intestinalization of the human gastric organoids following ectopic CDX2 expression ^28,60,61^.

In sum, these results confirmed that the ectopic expression of CagA in human gastric organoids leads to the activation of CDX2, a marker of GIM. Moreover, CagA-positive organoids acquired intestinal cell-like properties, such as downregulation of the stomach mucin Muc5AC and upregulation of intestinal mucin Muc2.

### Phosphorylation of C-term CagA is crucial for intestinal differentiation of human gastric organoids

The C-terminal region of CagA consists of a variable number of Glu-Pro-Ile-Tyr-Ala (EPIYA) motifs, which are tyrosine-phosphorylated upon CagA translocation ^11–13^. It has been observed that transgenic mice expressing wild-type CagA develop neoplasms in contrast to mice expressing a phosphorylation-resistant version of CagA, indicating a critical role for EPIYA phosphorylation *in vivo* ^18^. Thus, we evaluated if CagA-induced intestinal differentiation in gastric primary cells was dependent on the phosphorylation of the effector EPIYA. Using site-directed mutagenesis, we generated a phospho-resistant version of CagA (Fig. S3B and Fig. 1A) and transduced dissociated gastric organoids with a lentivirus expressing this mutant (ΔPR-CagA). Unlike organoids transduced with wild-type CagA, ΔPR-CagA-transgenic organoids did not show upregulation of CDX2 and MUC2 mRNA levels, supporting the theory that the upregulation of intestinal markers is CagA phosphorylation-dependent (Fig. 3F).

Finally, we sought to assess the impact of expressing only the C-terminal part of CagA in human gastric primary cells. For that purpose, dissociated gastric organoids were transduced with a construct containing a shorter version of CagA (C-term) (Fig. S3B and Fig. 1A). At 6 days post-transduction, the C-term CagA did not increase the CDX2 and MUC2 mRNA levels to the same degree as the full-length CagA (Fig. S3C). At 10 days of C-term CagA expression in gastric organoids, MUC2 mRNA levels were similar to those in gastric organoids expressing a full-length CagA; CDX2 expression remained lower (Fig. S3C). These results indicate that the C-terminal region of CagA is necessary but not sufficient to induce complete intestinal differentiation in human gastric organoids.

### The STAT3 pathway is activated in CagA-positive gastric organoids

Next, we wanted to determined which signaling pathways drive this CagA/CDX2-dependent intestinalization of human gastric organoids. From the transcriptional analysis data, STAT3 pathway target genes appeared to be upregulated, as shown by GSEA analysis (Fig. 4A). In support, previous studies have revealed that *H. pylori cag* PAI- and *vacA*-positive strains induce STAT3 activation in AGS and HEp2 cells and in Mongolian gerbils ^62,63^. Moreover, CagA-positive *H. pylori* human gastritis samples showed increased STAT3 activation ^36^, suggesting that STAT3 hyperactivation is driven by CagA. This prompted us to elucidate whether the expression of CagA activates the STAT3 pathway in human gastric organoids. We transduced single gastric primary cells with CagA lentivirus, allowing them to grow into 3-D organoid structures in expansion medium and harvested them at different time points. STAT3^Tyr705^ phosphorylation levels peaked at 6 days post-transduction compared to the non-transduced or *gfp-*lentivirus controls (Fig. 4B). CagA expression and its phosphorylation levels were maintained over at least seven passages (Fig. 4B). To evaluate STAT3 transcriptional activity, we examined the expression of the established STAT3-responsive gene SOCS3. A significant increase in SOCS3 mRNA was observed in concordance with STAT3 activation in CagA-positive gastric organoids at passage 1 (Fig. 4C).

**Fig. 4.**
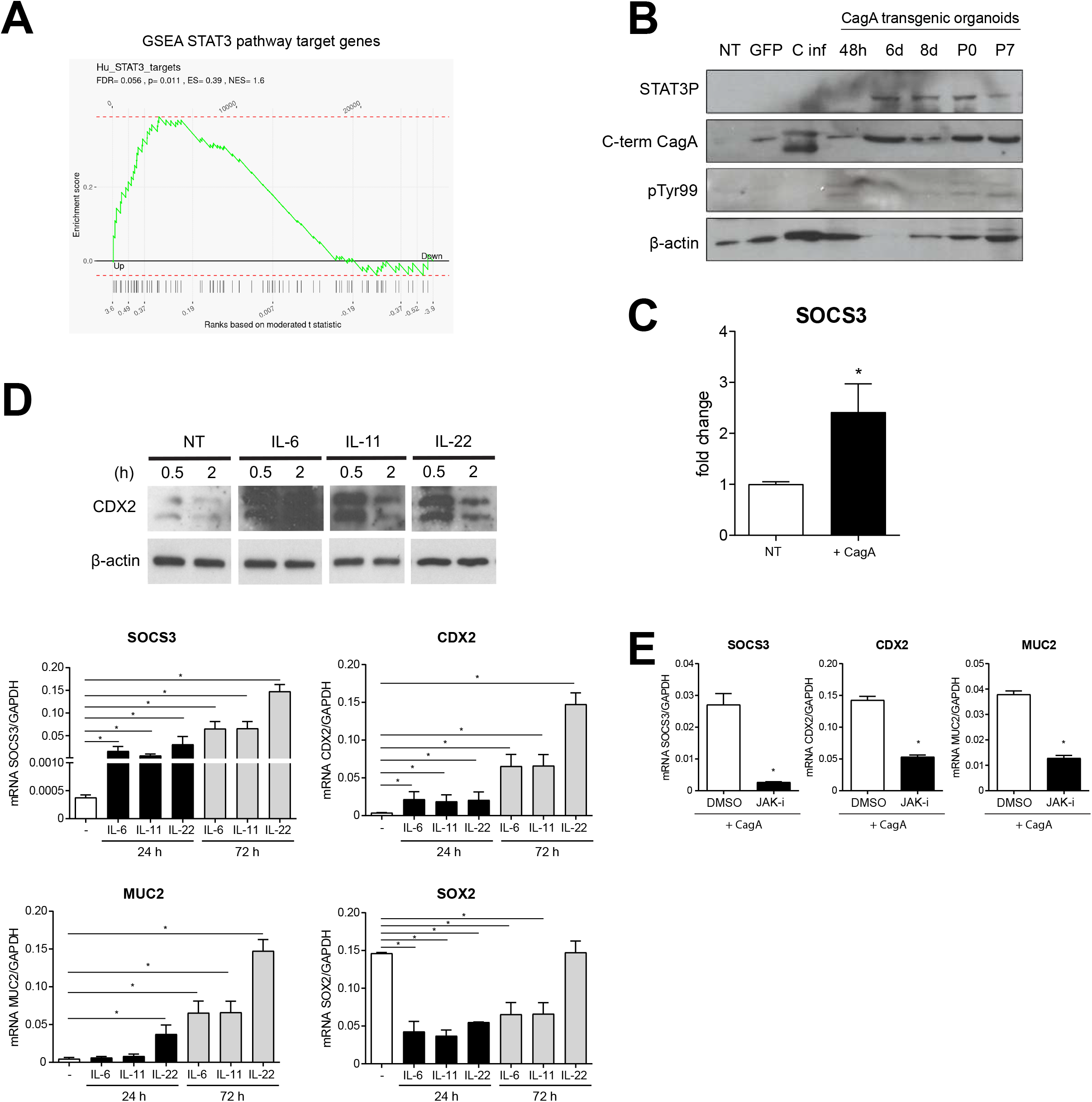
*H. pylori* CagA activates the STAT3 pathway in human gastric organoids leading to intestinal differentiation. (A) Gene set enrichment analysis of the microarray results based on the STAT3 target genes^85^. Genes are ordered according to their differential expression in CagA-positive organoids vs. control gastric organoids (left to right). STAT3 target genes are marked by black bars below the plot. The plotline indicates the running enrichment score. (B) Western blotting analysis of STAT3 activation in human gastric CagA positive organoids by detecting STAT3 phosphorylation: time points of 48 h, 6 and 8 days post-transduction, and at passages 0 and 7 after FACS. β-actin was used as a loading control. The results are representative of three independent experiments. (C) Gene expression analysis of SOCS3 by RT-qPCR in CagA-positive organoids at P0 after FACS. Values of mRNA levels were normalized against GAPDH expression and further normalized against non-transduced (NT) cells. Data represent mean ± SD of four independent biological experiments. (D) Western blotting analysis of CDX2 expression in healthy human gastric organoids upon treatment with IL-6 (100 ng/ml), IL-11 (100 ng/ml), and IL-22 (100 ng/ml) for 0.5 h or 2 h. Gene expression analysis of SOCS3, CDX2 and SOX2 RT-qPCR in healthy human gastric organoids upon treatment with IL-6 (100 ng/ml), IL-11 (100 ng/ml) and IL-22 (100 ng/ml) for 24 h or 72 h. Values of mRNA levels were normalized against GAPDH expression. (E) Gene expression analysis of SOCS3, CDX2 and MUC2 by RT-qPCR in CagA-positive organoids upon 18 h treatment with the STAT3 inhibitor S3BI100 (100μM) compared to DMSO as control. Values of mRNA levels were normalized against GAPDH expression.

IL-6 and IL-11, which can activate the JAK/STAT3 pathway, are the dominant IL-6 family cytokines expressed in the gastric mucosa ^37,64^. Furthermore, IL-22 has also been described to trigger STAT3 activation ^65^. Interestingly, some studies indicate that CDX2 can be regulated via IL-6 stimulation ^55^. To this end, we analyzed the association between STAT3 activation and CDX2 upregulation in human gastric organoids (Fig. 4D). Wild-type gastric organoids were treated with 100 ng/ml of IL-6, IL-11, and IL-22 at different time points and harvested for Western blotting and RNA extraction. Already after 0.5 and 2 h of treatment, gastric organoids expressed the CDX2 protein (Fig. 4D). STAT3 activation was also observed after 24 h and 72 h of treatment by all the cytokines, as shown by increased mRNA levels of its target gene SOCS3 (Fig. 4E). CDX2 mRNA expression also increased in each treatment group and at each time point (Fig. 4D), indicating that CDX2 expression depends on the activation of the STAT3 pathway. Furthermore, SOX2 expression was downregulated (Fig. 4D), confirming that the STAT3 activation negatively regulates SOX2 expression.

STAT3 activation by cytokines is mediated through the Janus family kinases (Jak) ^66^. Treating CagA-positive gastric organoids with a JAK inhibitor (AZD1480) ^67 68^, led to the downregulation of SOCS3, CDX2 and MUC2 mRNA levels, compared to controls treated with DMSO (Fig. 4F). This indicates that STAT3 activation is necessary for CDX2 overexpression in CagA-positive gastric organoids. Overall, our results show the involvement of the STAT3 pathway in CagA-induced intestinal trans-differentiation of human gastric primary cells.

## Discussion

Here, we established a transgenic gastric organoid model to elucidate the effects of the *H. pylori* CagA effector protein in normal human gastric epithelial cells. Strikingly, ectopic expression of CagA, occasionally referred as a bacterial oncoprotein ^16,69,70^, led to a marked phenotype of gastric intestinal metaplasia (GIM), which typically precedes the development of gastric cancer.

Expression of CagA in gastric organoids was followed by processing of the ∼135 kDa full-size protein to an EPIYA motif containing ∼35 kDa tyrosine-phosphorylated C-terminal fragment, termed p35^pTyr^. Processing of CagA has been described before in *H. pylori-*infected human cells lines, including primary professional phagocytes^44,46,71^, and appears to reflect a natural and efficient processing route in primary like-cells^47^. Previous results in well-studied transformed cell lines indicated via MALDI-MS analysis that the phosphorylated p35^pTyr^ corresponds to the C-terminal part of CagA^44^.

In this study, full-length CagA and the N-terminal product were not detected. This result was also obtained upon infection of AGS cells with *H. pylori* strains J99 and G27^45^. Similarly, the N-terminal product was not present in *H. pylori* cells grown without or decreased host cell contact^45,71^, comparably to our CagA transgenic organoids. In line with these studies, our results suggest that additional modifications occur in a *H. pylori* strain-specific manner and that the N-terminal could be a particular protein species of CagA modified in the host or sensitive to proteolysis. Several studies have described independent functions for the N-terminal and the C-terminal parts of CagA ^16^, but the mechanism responsible for processing remains obscure. The overall half-life of CagA upon translocation has been reported to be ∼200 min ^72^ and degradation may occur not via the proteasome but rather through autophagy^73^. Of note, we used CagA from a Western strain (P12) and, considering existing strain variations^45^, processing of CagA from East-Asian strains may differ.

Here we demonstrate trans-differentiation of the primary gastric epithelial cells to a phenotype that resembles intestinal cells as a response to CagA and CDX2 *de novo* expression. A similar phenotypic change has previously been seen in mice as the result of ectopic CDX2 expression in the stomach ^28,29^. Recently it was reported that in mouse gastric organoids the ectopic expression of CDX2 results in incomplete reprogramming towards intestinal-like cells ^59,74^. Strikingly, in our model CagA led to downregulation of gastric differentiation markers and enrichment of intestine-specific genes and GIM markers, as specified by Companioni et al ^48^, indicating that activated CagA leads to trans-differentiation of adult gastric stem cells towards intestinal-like stem cells. Moreover, ectopic expression of CDX2 in the gastric organoids also led to downregulation of SOX2. This observation supports a negative regulation of the stomach-specific marker SOX2 by CDX2 ^55 51,74–76^. Of note, downregulation of SOX2 has been also observed both in the *H. pylori-*infected stomach and GIM ^51^.

Moreover, we demonstrate with CagA-transgenic gastric organoids that CDX2 *de novo* synthesis is a downstream effect of CagA-dependent STAT3 signaling activation. While the mechanism of CagA dependent STAT3 activation remains to be elucidated, it has previously been suggested that CagA indirectly regulates the levels of phosphorylated STAT3 by interacting and sequestering SHP2, the major negative regulator of STAT3 ^77^. Other pathways may also be involved in the regulation of CDX2 expression. For example, it has been shown that BMP signaling is activivated in GIM, co-localising with CDX2 expression^78^; it would be of interest to confirm this in primary cells.

Our model of human gastric organoids may recapitulate tissue-specific responses^36^ more accurately than previous studies using transformed cell lines. As determined by cytokine treatment of gastric organoids, we found that IL-11, IL-6 and IL-22 are able to activate the STAT3 pathway. An increase of IL-11 expression is observed in CagA-expressing MKN28 gastric cells ^79^. Although we did not observe significant increase of cytokine or respective receptor expression in our arrays, an assessment of cytokine synthesis and secretion is still pending. Inhibition of JAK2, which is downstream of the gp130 cytokine receptor and upstream to STAT3 phosphorylation ^37^, caused a reversion of the expression of intestinal markers in CagA-transgenic organoids supporting the significance of STAT3 signalling in the initiation of GIM^38^.

Along with increased CDX2 expression, we noticed upregulation of the intestinal mucin MUC2 and downregulation of the stomach mucin MUC5AC. These changes in mucin expression could be mirrored by ectopically expressing CDX2 in the gastric organoids, emphasizing the role of CDX2 as a master controller of intestinal trans-differentiation of adult gastric stem cells. The mucin profile described in this study could also be related to STAT3. Likewise, the expression of MUC4 was shown to be dependent on STAT3 activation ^80^. Other studies have identified a link between CDX2 and claudin-2 activation mediated through STAT3 phosphorylation ^55,57^. Moreover, we observed in our arrays an increased expression of REG1A and REG3A, potentially downstream of STAT3 activation. These bactericidal lectins might provide *H. pylori* a competitive advantage by selectively reducing the fitness of, e.g. Gram-positive, bacteria present in the microbiota of the gastric mucosa in favor of its own survival ^77^. Interestingly, human β-defensin 3 (hBD3), a highly effective defensin against *H. pylori* ^81–83^, is slightly down-regulated in the CagA-positive gastric organoids. We have previously shown that *H. pylori* prevents hBD3 expression through a CagA-dependent mechanism^83^.

In summary, we established a human gastric organoid model to study *H. pylori*-CagA effects on gastric cell differentiation and possible cellular transformation events. Our work provides molecular insights into the role of CagA in the pathological cascade of intestinal-type gastric carcinoma development. This bacterial effector protein promotes intestinal differentiation of human gastric epithelial cells through the aberrant expression of CDX2 via STAT3 pathway activation. To this end, these findings advance the understanding of the initial processes of GIM and cancer development in the human stomach.

## Materials and Methods

### Ethical permissions

Human gastric tissue samples for the preparation of primary cells were derived from the Clinics for General, Visceral and Transplant Surgery, and the Center of Bariatric and Metabolic Surgery, Charité University Medicine, Berlin, Germany. Scientific usage of the samples for experimental purposes was approved by the ethics committee of the Charité University Medicine, Berlin (EA1/058/11 and EA1/129/12).

### Human gastric primary cells culture and media

Pseudonymized samples (see Table S1) were obtained from individuals undergoing gastrectomy or sleeve resection and processed as described previously by our group for organoid cultures ^39^.

### Construction of lentivirus plasmids

To generate CagA expression vectors, full-length *cagA*-3647bp cDNA from P12 *H. pylori* strain (strain collection number 243, EPIYA motifs ABCC) was amplified by PCR using Phusion polymerase, adding *KpnI* and *XhoI* restriction sites with matching primers. After A’ tailing, the PCR product was cloned into pGEMTeasy (Promega A1380). The *cagA* fragment was digested with *KpnI* and *EagI* and subcloned into the lentivirus entry vector pEF-1α/pENTR (Addgene, #17427) containing the EF promoter. The *cagA* cDNA with the EF promoter was recombined *in vitro* with the viral Destination vector pLenti CMV GFP DEST (Addgene, #19732) using the Gateway System (Invitrogen). Prior to lentivirus preparation, sequencing of each vector containing *cagA* and transfection of the expression vector in AGS cells was performed to confirm the correct sequence of *cagA* and verify the correct expression of CagA (data not shown).

To generate HA-tagged constructs, pEF_*cagA* constructs were used as a template. We designed PCR primers: reverse containing SphI restriction site and forward containing the HA-tag, STOP codon and *XhoI* restriction site. We performed a PCR that generated a 300bp product, purified it, digested it with *SphI* and *XhoI* and cloned it into pEF-*cagA,* digested by *SphI* and *XhoI*. As explained before, *cagA*-HA constructs were recombined using Gateway Cloning into the destination vector pLentiCMV GFP DEST (Addgene, #19732). Sequence and correct expression of CagA-HA were verified (data not shown).

To generate phospho-resistant *cagA* constructs, pEF_*cagA* constructs were used as a template for site-directed mutagenesis, precisely mutating four tyrosine sites present in the EPIYA motifs ^43^ (Fig. S3B). Briefly, a PCR of the C-term region of *cagA* was performed flanking the fragment with *MunI* and *SphI* restriction sites. After A’ tailing, the PCR product was cloned into pGEMTeasy (Promega A1380). The mutagenesis of the EPIYA sites were done one after the other using the kit Quick change II site-directed mutagenesis (Aligent #200523). The primers to mutate each motif are listed in Table S2. The resulting pEF plasmid, which contained full *cagA* with the four EPIYA motifs mutated to EPIAA, was recombined into the lentivirus destination vectors as explained before.

To generate a shorter version of *cagA,* pEF_*cagA* constructs were used as a template. The plasmid was digested with KpnI and SwaI cutting out the N-terminal part of CagA. The fragment ends were filled in with T4 polymerase and dNTPs and ligated into the corresponding pLenti vectors. The resulting fragment coded a shorter protein of 67 Kda CagA C-term (from 1629 bp to 3647 bp) (Fig. S3B).

CDX2-overexpressing constructs were designed and purchased online using Vector Builder. The plasmid construct is illustrated in Fig. S3A. Briefly, *cdx2* expression is under the control of CAG promoter and coupled with a 2A sequence to the successive expression of *m-cherry*.

### Lentiviral manipulation

Replication-deficient lentiviral particles were produced by CaCl2-transfection of 293-T cells with the packaging vector psPAX2 (Addgene, #12260), the envelope vector pMD2.G (Addgene, #12259) and pLVTHM (Addgene, #12247). After two days, the supernatant containing the produced lentiviral particles was filtered (0.45 μm), concentrated with Lenti-X Concentrator (Clontech), and the pellet was dissolved in ADF medium (Thermo Fischer #12634). Organoid cultures were prepared as described for passaging ^39^ and collected in 500 μl infection medium/sample (expansion medium with ADF containing lentiviral particles instead of normal ADF plus 10 μg/ml polybrene (Sigma)). The cell suspension was transferred to a 24-well plate coated with 30 µl of Matrigel (growth factor reduced, phenol red-free; BD Biosciences). After 24 h, the medium containing the lentivirus and the non-attached primary cells was removed. The attached primary cells were washed twice with PBS and covered with 50 µl of Matrigel. After 15 min at 37°C, 500 µl of expansion media was added, and the organoids were cultured under normal conditions ^39^.

### FACS enrichment of transgenic primary gastric epithelial single cells

Organoid cultures were prepared as described for passaging ^39^ and collected in 500 µl of sorting buffer (PBS, 1% FCS, 1% HEPES, 1% DNAse). Single cells were filtered through 40 µm filters into a polystyrene FACS tube. GFP-positive single cells were sorted by FACS with the exclusion of dead cells using propidium iodide. Sorted single cells were collected in ADF plus 10% FCS and 9 µM Rock inhibitor (Y-27632, Sigma) to prevent anoikis. An amount of 5 x 10^5^ single cells was embedded in Matrigel, seeded in 24 well-plate, and maintained as described^39^.

### Cytokine treatment in 3D organoids

Gastric organoids were grown as described ^39^ and the cytokine treatments were done in 3D. Recombinant human IL-6, IL-11 and IL-22 (Prepotech 100 ng/ml) were added to selected wells and maintained at indicated times until the end of the experiment.

### Protein lysates and immunoblot analysis

Cells were directly harvested with 2x Laemmli buffer (4% SDS, 29% glycerol, 120 mM Tris-Cl (pH 6.8) and 0.02% bromphenol blue) and boiled for 10 min at 95°C. Samples were separated on a 10% SDS-PAGE-polyacrylamide gel and transferred to PVDF membranes by standard procedures. We used antibodies against N-terminal CagA b-300 (Santa Cruz Cat. No. sc-25766), C-terminal CagA bk-20 (Santa Cruz Cat. No. sc-48128), p-Tyr (Santa Cruz Cat. No. sc-7020), HA (Santa Cruz Cat. No.sc-805), pY705STAT3 (Cell Signaling, Cat. No. #4113), CDX2 [EPR2764Y] (Abcam Cat. No. ab76541), Claudin-2 (Abcam Cat. No. ab53032), β-Actin (Sigma, Cat. No. A5441), and horseradish peroxidase (HRP)-conjugated anti-mouse, anti-rabbit or anti-goat secondary antibodies (Amersham). In all cases, the signal was detected with the WESTERN LIGHTNING™ western blot kit system for ECL immunostaining. Antibodies used are listed in Table S4.

### RT-qPCR

RNA extraction was performed using the GeneJet RNA purification kit (Fermentas) according to the manufacturer’s instructions. Conversion of RNA into cDNA and subsequent cDNA amplification was achieved together by using Power Sybr^®^ Green RNA-to-CT™ 1-step kit (Applied Biosystems) and 7500 fast real-time PCR system (Applied Biosystems) following the manufacturer’s instructions. Primer sequences are listed in Table S3.

### Immunofluorescence staining

Organoids were prepared using the same protocol ^39^ for passaging, after which they were collected in 2D medium and seeded in collagen-coated 24 well-plate containing glass coverslips. Cultures were kept in a humidified incubator at 37°C, 5% CO2. Primary 2D cell cultures were grown to a confluency of 80%, washed twice with PBS, and fixed with 3.7% paraformaldehyde (Sigma) for 20 min. Primary antibodies and dyes (E-cadherin BD Bioscience #610181, dilution 1:100; CDX2 [EPR2764Y] Abcam ab76541, dilution 1:100; Hoechst Sigma H6024, dilution 1:10000) were diluted in blocking solution (0.3% BSA, 10% FCS, and 0.3% Triton in 1X PBS) and incubated overnight at 4 °C. Secondary antibodies were diluted in a blocking solution and incubated for 45 min at room temperature. Mounting was done with Mowiol (Vectorlabs, Inc.). Images were acquired with a Leica TCS SP-8 confocal microscope and processed using ImageJ 1.52. Antibodies used are listed in Table S4.

### Microarray expression profiling and data analysis

Total RNA was isolated with TRIzol (Life Technologies) according to the supplier’s protocol. RNA was labeled and hybridized on Agilent Human Gene Expression v2 4×44K microarrays or Agilent custom human whole-genome 8×60k arrays, using single-color hybridization according to the manufacturer’s instructions. Per-probe intensities were extracted using Agilent Feature Extraction software. Data were background corrected and normalized using the R package LIMMA ^84^, which was also applied to determine differential gene expression. Gene set enrichment analysis (GSEA) was performed on combined log2 fold changes from both platforms using R package fgsea with standard settings and 5000 permutations. The gene signature of STAT3 activation was derived from Hu et al. ^85^. The GIM signature was derived from published microarray data ^48^ in GSE78523 as genes upregulated with log2 fold change greater than 1 and adjusted p-value less than 0.05. Small intestine-specific genes were obtained from Protein Atlas (www.proteinatlas.org). Genes corresponding to the signatures of Lgr5 stem cells were taken from Suppl. Table 3 in Munoz et al. ^86^. The colon vs. stomach signature was generated by using differentially expressed genes (log2FC > 1.5, FDR < 5%) from a comparison of colon biobank normal tissues ^87^ and gastric glands ^41^. CDX2 target genes were taken from Boyd et al ^58^.

Microarray data will be deposited in the Gene Expression Omnibus (GEO; www.ncbi.nlm.nih.gov/geo/) of the National Center for Biotechnology Information and can be accessed with the GEO accession number GSE246474.

### Statistical Analysis

Statistical significance was analyzed using paired two-tailed Student’s t-test.

## Acknowledgments

The authors would like to thank Dr. Kfir Lapid and Dr. Rike Zietlow for editing the manuscript, Jörg Angermann for technical assistance in lentivirus production, and Diane Schad for help generating the graphics.

## Figure legends

**Supplementary figure 1.**
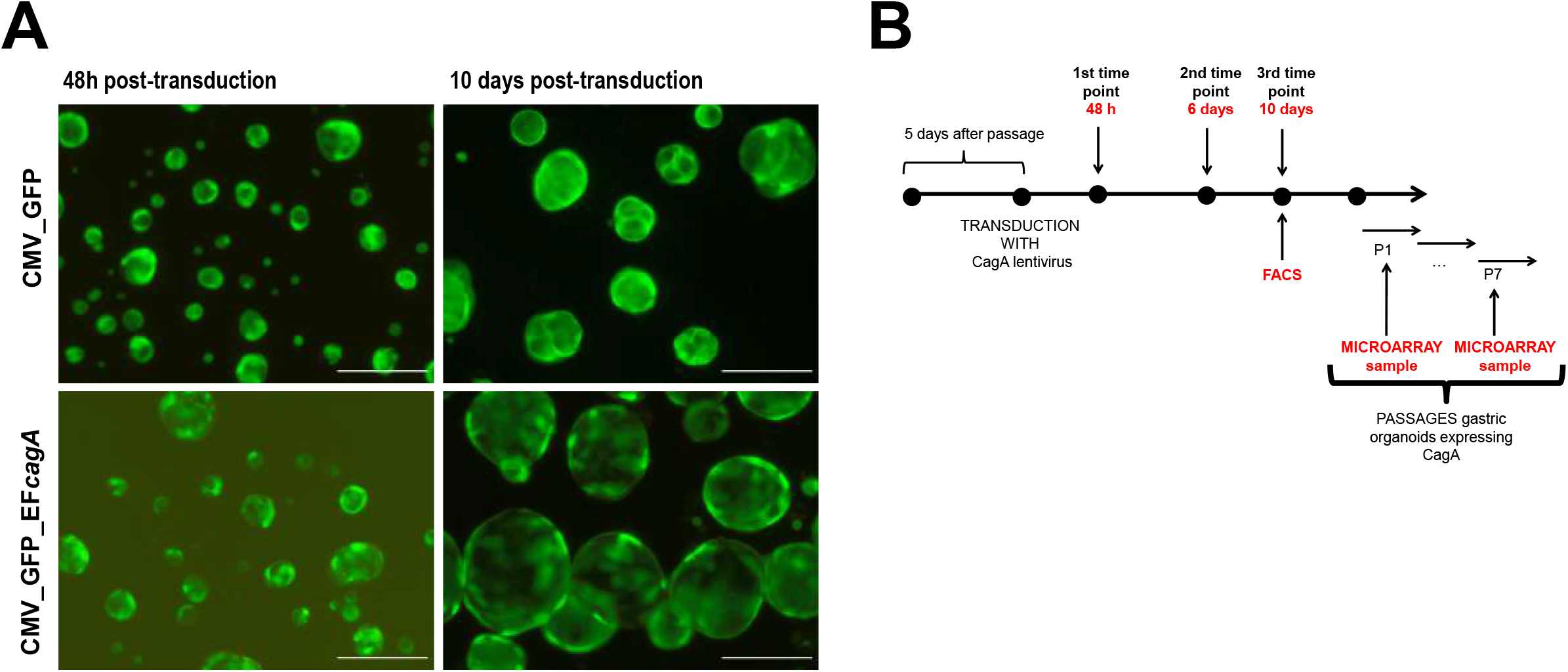
(A) Representative micrographs of human gastric organoids transduced with a lentivirus containing CMV_GFP_EF_*cagA* and CMV_GFP at 48h and 10 dayspost-transduction. Scale bars: 500µm. (B) The scheme describes the strategy to generate CagA-positive human gastric organoids using the lentivirus transduction, indicating the time points used for RT-qPCR, WD and microarray analysis.

**Supplementary figure 2.**
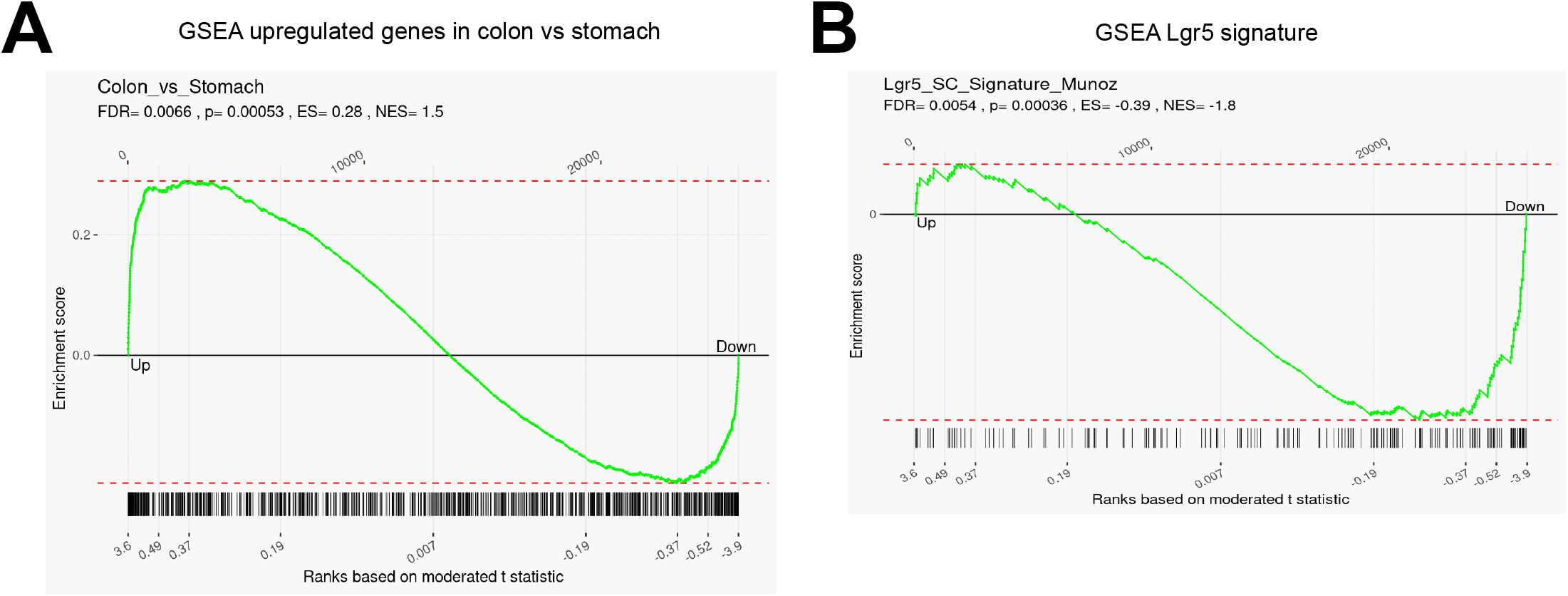
(A) Gene set enrichment analysis of the microarray results based on genes enriched in the colon as compared to the stomach. Genes are ordered according to their differential expression in CagA-positive organoids vs. control gastric organoids (left to right). Upregulated genes in the colon tissue vs. the stomach tissue are marked by black bars below the plot. The plotline indicates the running enrichment score. (B) Gene set enrichment analysis of the microarray results based on Lgr5 target genes^86^. Genes are ordered according to their differential expression in CagA-positive organoids vs. control gastric organoids (left to right). Intestinal stem cell genes are marked by black bars below the plot. The plotline indicates the running enrichment score.

**Supplementary figure 3.**
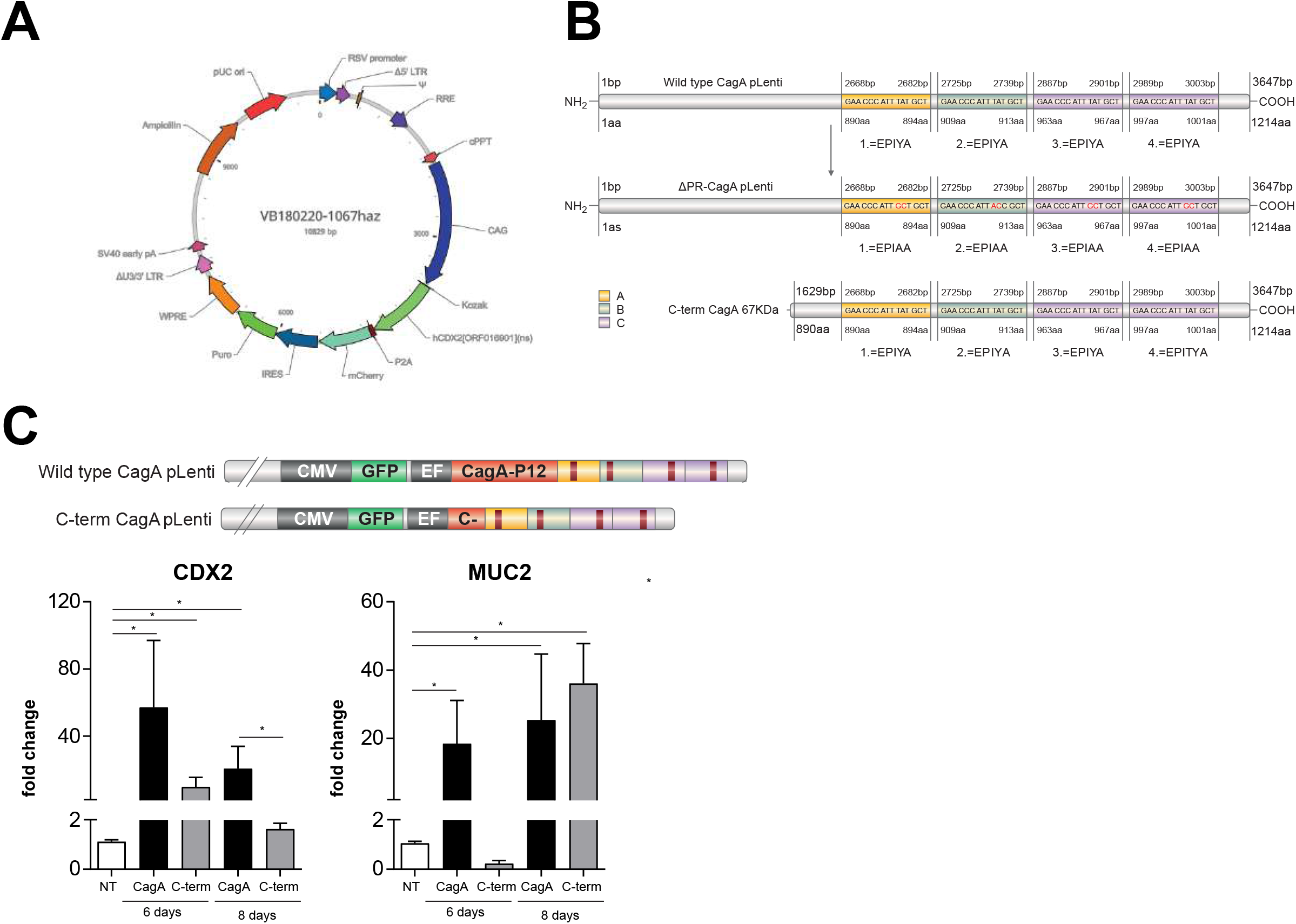
(A) The lentivirus vector was designed using Vector Builder to overexpress CDX2 in gastric organoids. Briefly, *cdx2* expression is under the control of the CAG promoter and coupled with a 2A sequence for the consequent expression of *m-cherry*. (B) Schematic presentation of the site-directed mutagenesis in wild-type CagA phosphorylation sites. Single phenylalanine substitutions of tyrosine residues in phosphorylation motifs were introduced into constructs. Schematic presentation of the C-terminal CagA fragment of 67KDa cloned in the lentivirus. (C) Analysis of CDX2 and MUC2 expression by RT-qPCR in CagA-positive organoids (CagA) and in organoids expressing a shorter mutant CagA (C-term). Values were normalized against GAPDH expression. Data show mean ± SD of three independent biological experiments.

## Notes

**Funding:** This work was supported by the European Research Council (ERC) Advanced Grant (885008-MADMICS) to T.F.M. and the EMBO long-term fellowship ALTF 580-2014 to M.R.

### Competing Interest Statement

The authors have declared no competing interest.

