## supplementary tables for "Helicobacter pylori cancer associated CagA protein drives intestinal metaplastic transition in human gastric organoids"

Supplementary table 1

Patients samples used for the study

| ID | Part | Age | Sex | Comments |
|---|---|---|---|---|
| GAT15 | Antrum | 47 | Male | BMI:52, Hpy Diabetes |
| GAT16 | Antrum | 34 | Male | BMI:56, Hpy Negative |
| GAT18* | Antrum | 50 | Female | BMI:43, Hpy Negative |
| GAT23A* | Antrum | 55 | Female | BMI:45, Hpy Negative (Dennecke) |

*Used for microarray analysis

Supplementary table 2

Primers used for site-directed mutagenesis and C-term cloning of CagA

| **Primer** | **Sequence (5' -> 3')** |
|---|---|
| EPIYA 1 Fw | GACTCAAAAACGAACCCATTTATGCTAAAG |
| EPIYA 1 Rv | ATTAACTTTAGCATAAATGGGTTCG |
| EPIYA 2 Fw | GCCATGAAGAACCCATTTACGCTCAAG |
| EPIYA 2 Rv | GCAACTTGAGCGTAAATGGGTTCTTC |
| EPIYA 3+4 Fv | AGCTAGCCCTGAACCCATTTATGCTACG |
| EPIYA 3+4 Fv | GTAGCATAAATGGGTTCAGGGC |
| C-term Fw | AGCAAGCGTTAGCCGATCTC |
| C-term Rv | CACGAGCTTGAGCCACTCAG |

Supplementary table 3

RT-PCR primers

| | |
|---|---|
| CagA P12 | Sequence (5' -> 3') |
| Forward Primer | AACAACCACAAACCGAAGCG |
| Reverse Primer | ATCGTATGAAGCGACAGCGT |
| hMUC6 | Sequence (5' -> 3') |
| Forward Primer | CAGCTCAACAAGGTGTGTGC |
| Reverse Primer | TGGGGAAAGGTCTCCTCGTA |
| hMUC5AC | Sequence (5' -> 3') |
| Forward Primer | GGAGGTGCCCACTTCTCAAC |
| Reverse Primer | CTTCAGGCAGGTCTCGCTG |
| hSOX2 | Sequence (5' -> 3') |
| Forward Primer | TACAGCATGTCCTACTCGCAG |
| Reverse Primer | GAGGAAGAGGTAACCACAGGG |
| hCDX2 | Sequence (5' -> 3') |
| Forward Primer | CGGCAGCCAAGTGAAAAC |
| Reverse Primer | CGGATGGTGATGTAGCGACT |
| hMUC2 | Sequence (5' -> 3') |
| Forward Primer | GAGGGCAGAACCCGAAACC |
| Reverse Primer | GGCGAAGTTGTAGTCGCAGAC |
| hSOCS3 | Sequence (5' -> 3') |
| Forward Primer | GGAGACTTCGATTCGGGACC |
| Reverse Primer | GAAACTTGCTGTGGGTGACC |
| GAPDH | Sequence (5' -> 3') |
| Forward Primer | GGTATCGTGGAAGGACTCATGAC |
| Reverse Primer | ATGCCAGTGAGCTTCCCGTTCAG |

Supplementary table 4

Antobodies and dies

| Antibody | Supplier | Cat. No | dilution |
|---|---|---|---|
| CDX2 | Abcam | ab76542 | WB 1:500<br>IF 1:100 |
| Claudin-2 | Abcam | ab53032 | WB 1:1000 |
| bk-20 C-term CagA Ab | Santa Cruz | sc-48128 | WB 1:1000 |
| m-300 N-term CagA Ab | Santa Cruz | sc-25766 | WB 1:1000 |
| p-Tyr | Santa Cruz | sc-7020 | WB 1:1000 |
| pY705STAT3 | Cell Signaling | 4113 | WB 1:1000 |
| β-Actin | Sigma | A5441 | WB 1:10000 |
| Hoescht | Sigma | H6024 | IF 1:10000 |
